# AGFusion: annotate and visualize gene fusions

**DOI:** 10.1101/080903

**Authors:** Charlie Murphy, Olivier Elemento

## Abstract

**Summary:** The discovery of novel gene fusions in tumor samples has rapidly accelerated with the rise of next-generation sequencing. A growing number of tools enable discovery of gene fusions from RNA-seq data. However it is likely that not all gene fusions are driving tumors. Assessing the potential functional consequences of a fusion is critical to understand their driver role. It is also challenging as gene fusions are described by chromosomal breakpoint coordinates that need to be translated into an actual amino acid fusion sequence and predicted domain architecture of the fusion proteins. Currently there are no easy-to-use tools that can automatically reconstruct and visualize fusion proteins from genomic breakpoints. To facilitate the functional interpretation of gene fusions, we developed AGFusion, available as an online web tool that can be readily used by non-computational researchers as well as a python package that can be built into computational pipelines. With minimal input from the user, AGFusion predicts the cDNA, CDS, and protein sequences of all gene fusion products based on all combinations of gene isoforms. For protein coding fusions, AGFusion can annotate and visualize the protein domain architecture. AGFusion currently supports *Homo sapiens* (genome builds GRCh37 and GRCh38) and *Mus musculus* (genome build GRCm38) and new genomes can easily be added.

**Availability:** AGFusion python package is freely available at https://github.com/murphycj/AGFusion under the MIT license. The AGFusion web app is available at http://agfusion.info

## 1 Introduction

Gene fusions are the result of structural chromosomal rearrangements and can cause a variety of functional changes for the involved genes. Frequently, gene fusions can produce a chimeric protein, whereby the protein domains are combined in a way to produce novel functions. Gene fusions can also result in the rearrangement of regulatory regions such as promoter elements and 3' or 5' UTRs, thereby altering the expression of the involved genes. The advent in next-generation sequencing has revealed a large and heterogeneous number of gene fusions in many different types of cancer ^1,2^. The functional role of many fusions has been elucidated, but certain cancers such as breast cancer have presented with a diverse array of gene fusions and understanding their function and role in driving tumors or therapy resistance is critical. This is important as many gene fusions are therapeutically targetable and will likely play a growing role in precision medicine ^2,3^.

Several software packages for gene fusion visualization exist, each specializing in displaying certain information about a gene fusion. These tools have significant limitations. Both iFUSE and Transcriptome Viewer (TViewer) can predict the DNA, RNA, and protein sequences from the fusion as well as visualize the fusion gene and transcript models, respectively—but neither annotate or visualize the resulting protein products ^4,5^. The Integrative Genomics Viewer (IGV) can display the fusion-supported mapped reads and also allows a split view mode to visualize reads mapping to two disparate genomic locations ^6^. The DEXSeq R package can allow the user to examine expression pattern across exons, which is useful to confirm expression changes in individual exons before and after the fusion junction ^7^. Circos plots can display gene fusions as arcs between genomic locations, which are arranged in a circular manner ^8^. Pegasus is a computational tool that can predict and annotate gene fusion protein sequences and assign an oncogenic score, but provides no visualization ^9^. However, to the best of our knowledge, there is no fully integrated software that predicts DNA, RNA, and protein sequences but also annotates and visualizes the resulting protein domain architecture of a gene fusion while taking into consideration the complexity of gene isoform combinations. To address this gap, we developed AGFusion as a web tool for non-computational users as well as a Python package that can be easily integrated into computational pipelines.

## 2 Implementation

Basic usage of AGFusion needs as input the 5' and 3' gene partners and their predicted junction locations. Currently, AGFusion only allows inputting of gene fusions that are in the coordinates of the human genome assemblies GRCh38 and GRCh37 and mouse genome assembly GRCm38. The user can input either the Ensembl Gene ID or the HGNC gene symbol or MGI gene symbol for the human or mouse genomes, respectively. AGFusion predicts the effect of the gene fusion for all gene isoform combinations. It can output the predicted full-length cDNA sequences, and if the fusion produces a protein product, the CDS and protein sequences are predicted as well as annotated with Pfam domains ^10^. Publication quality figures of the protein domain architecture can then be saved (Figure 1). Customization features allow the user to provide different names and colors for the protein domains.

**Figure 1:**
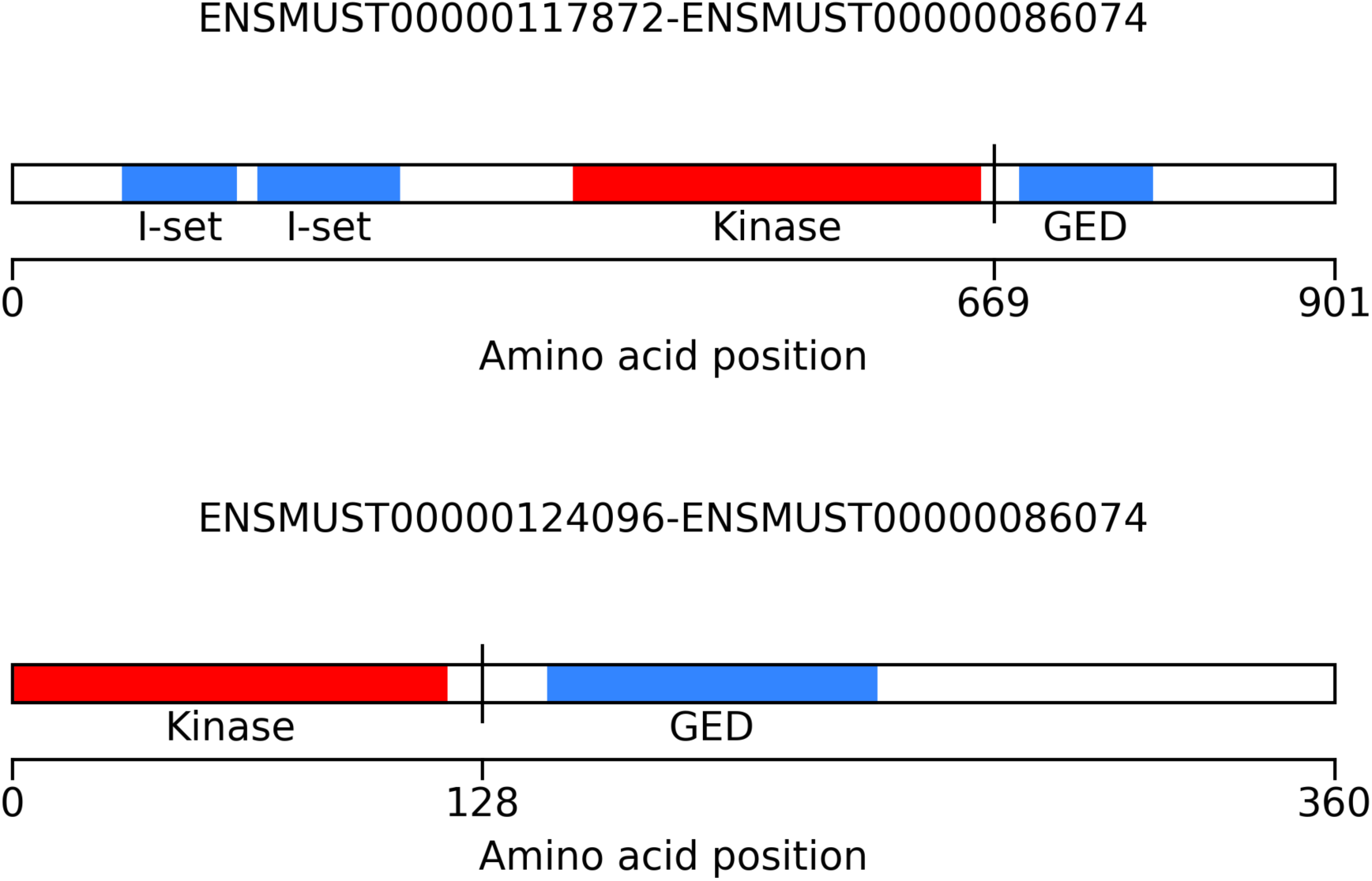
This example visualization shows two different isoforms of an FGFR2-DNM3 in-frame fusion found in a mouse tumor. The FGFR2 and DNM3 gene isoform combinations are shown at the top of each figure and the fusion junction indicated by a vertical line at amino acid positions 669 (top) and 128 (bottom). Each isoform contains intact kinase and GED domains, but they differ in I-set domains—indicating how important it can be to consider gene isoforms combinations that form a fusion.

AGFusion depends on data from Ensembl and Pfam to annotate and visualize any user-inputted gene fusion ^10,11^. Ensembl data is provided primarily by the PyEnsembl python package, but the Biomart python package is used to download protein domain annotations for each Ensembl transcript ID ^12^. The Pfam data is then stored in a simple SQLite database that comes with AGFusion for faster querying. The SQLite database and PyEnsembl allow the AGFusion python package to be used without the need for an Internet connection.

The AGFusion web application is built as a graphical interface to the AGFusion python package for non-computational users. After the user inputs the gene fusion information the web app outputs figures that visualize the protein domain architecture of all fusion isoforms. The images are exportable as PDF or PNG format. The user can also download FASTA files of the fusion cDNA, CDS, and protein sequences.

## 3 Conclusion

Gene fusions are a heterogeneous source of tumor drivers and AGFusion can speed up the process of functional elucidation by annotating and visualizing the fusion product. Both computational and non-computational researchers can easily use AGFusion. We plan on expanding the function of AGFusion in several directions. Besides adding more customization and usability features, we plan on including all protein domain annotation used in Ensembl as well as adding support for more human and mouse genome assemblies. We also plan on adding a REST API to the AGFusion website.

## Funding

C.M. was supported by the Tri-Institutional Training Program in Computational Biology and Medicine via NIH training grant 1T32GM083937. O.E. was supported by NIH grants R01CA194547 and.

